# Real-world body orientation impacts virtual navigation experience and performance

**DOI:** 10.1101/2023.02.10.527966

**Authors:** Hyuk-June Moon, Hsin-Ping Wu, Emanuela De Falco, Olaf Blanke

## Abstract

Most human navigation studies in MRI rely on virtual navigation. However, the necessary supine position in MRI makes it fundamentally different from daily ecological navigation. Nonetheless, until now, no study has assessed whether differences in physical body orientation (BO) affect participants’ experienced BO during virtual navigation. Here, combining an immersive virtual reality (VR) navigation task with subjective BO measures and implicit behavioral measures, we demonstrate that physical BO (either standing or supine) modulated experienced BO. Also, we show that standing upright BO is preferred during spatial navigation: participants were more likely to experience a standing BO and were better at spatial navigation when standing upright. Importantly, we report that showing a supine virtual agent reduces the conflict between the preferred BO and physical supine BO. Our study provides critical, but missing, information regarding experienced BO during virtual navigation, which should be considered cautiously when designing navigation studies, especially in MRI.

**Visual Abstract:** 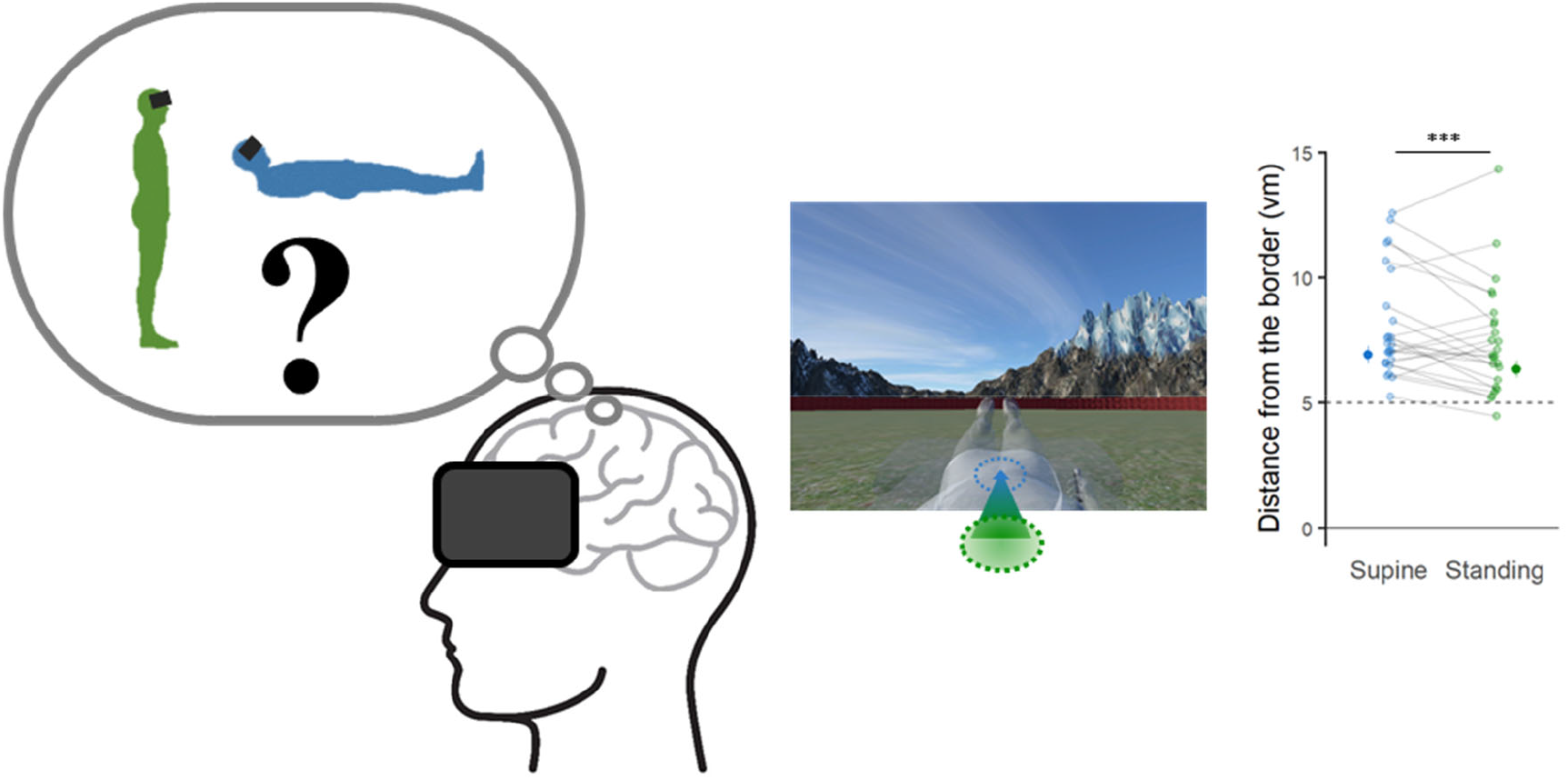

## Introduction

The brain mechanisms of spatial navigation in humans are a prominent topic in the basic neurosciences (Maguire et al., 1999; Buzsaki and Moser, 2013; Ekstrom and Isham, 2017; Bellmund et al., 2018) and are of clinical relevance (deIpolyi et al., 2007; Coughlan et al., 2018). The vast majority of human spatial navigation studies have used virtual navigation paradigms due to the fact that most of the non-invasive brain imaging techniques do not allow subjects to navigate in the real world and require participants to remain immobile. One of the most frequently used brain imaging techniques for human spatial navigation research, for example, is functional magnetic resonance imaging (fMRI). While fMRI provides access to neural activities in the deep brain structures, including the medial temporal lobe that are known to be crucial for spatial navigation (Scoville and Milner, 1957; Byrne et al., 2007; Moser et al., 2008; Squire, 2009), it requires participants not only to remain as immobile as possible, but also to be in a supine position. Although previous virtual navigation in fMRI significantly contributed to our understanding of human spatial navigation systems, the supine position in the MRI scanner imposes a fundamental difference between virtual navigation “in the scanner” and ecological daily navigation “in the real world”. However, still, how these differences impact human navigation systems still needs further investigation (Taube et al., 2013; Park et al., 2018; Steel et al., 2021).

Thus, it is unknown whether and how differences in body orientation (BO) of the physical body affect (1) subjective experience of BO during virtual navigation (i.e., what is the experienced BO during virtual navigation when subjects are in a supine physical BO?) and (2) spatial navigation performance (e.g., navigation accuracy or speed). To the best of our knowledge, no study has addressed this issue directly. Also, it is often assumed that, regardless of their physical BO, participants in navigation studies experience themselves as if they were standing upright during virtual navigation(Jacobs et al., 2013; Taube et al., 2013; Maidenbaum et al., 2018). However, as suggested by Moon and colleagues(Moon et al., 2022), bodily signals (e.g., vestibular and proprioceptive signals) from the physical body (i.e., supine participants in the scanner) can affect how participants experience the virtual agent in a VR environment. The bodily reference frame of the BO of the participant may thus be in conflict with the bodily reference frame of the virtual agent’s BO (and such mechanism may even be at play when no navigating avatar is shown in the virtual environment, as done in most previous spatial navigation research). In the present study, we hypothesized that the subjectively experienced BO in the virtual navigation space (as well as spatial navigation behavior) depends on the participant’s physical BO (in the scanner; mediated through intrinsic bodily signals) and, additionally, whether an avatar is shown in VR or not. To test this hypothesis, we asked participants to perform the same, classical, spatial navigation task (Doeller et al., 2010) in two different physical BOs (either supine or standing). Based on findings by Moon et al. (2022), we also tested two additional conditions where a supine virtual agent was shown or not (see Fig.1a and Methods).

**Figure 1.**
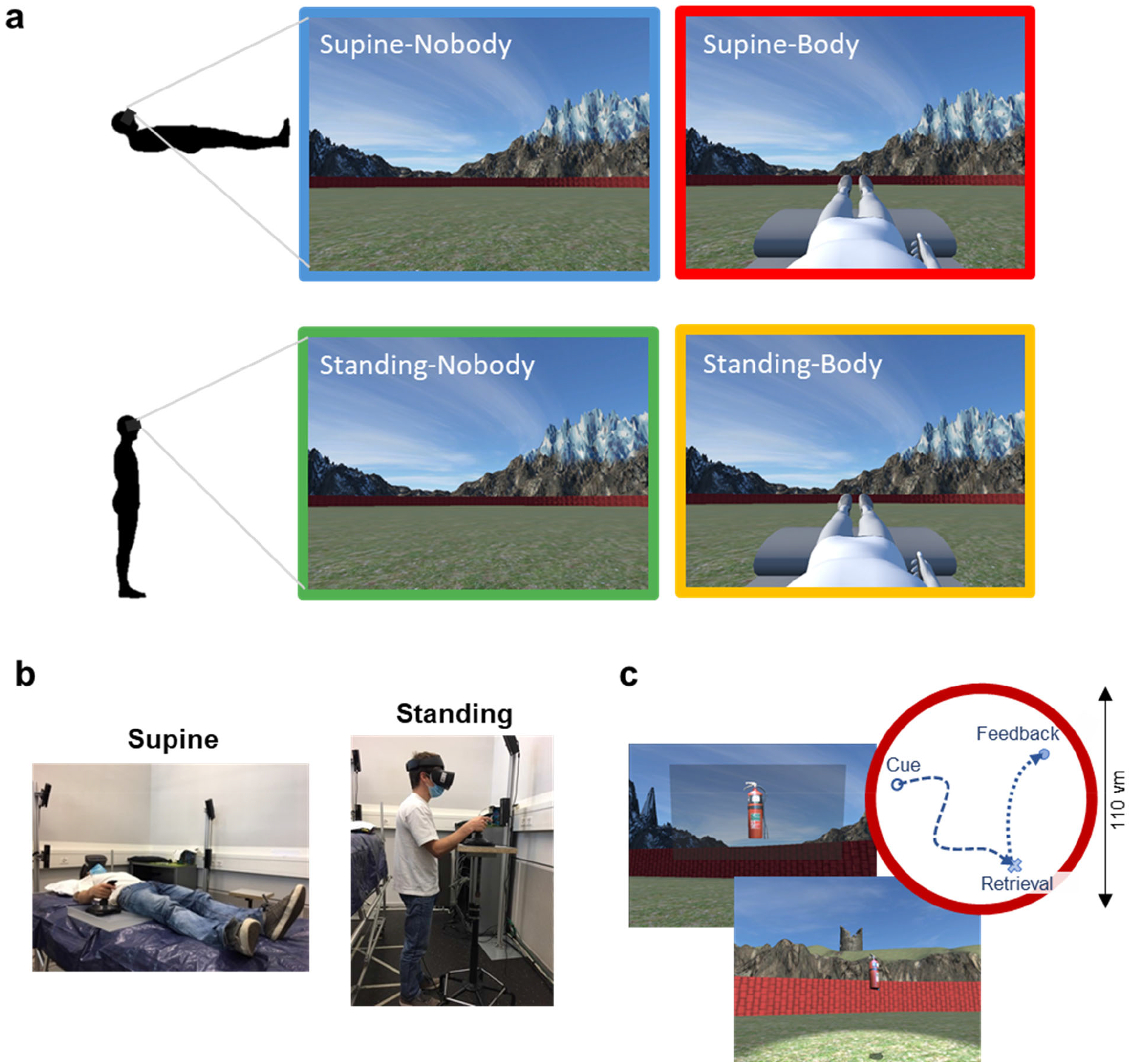
Experimental design combining two physical body orientations (BOs) and the presence/absence of an avatar conditions. **a,** Participants performed the task in four different conditions obtained by the combination of the physical body orientation (BO) (i.e., Supine vs. Standing) and the presence/absence of the avatar (i.e., Body vs. Nobody). Note that the avatar (when shown) was always in a supine position. **b,** Participants performed the virtual navigation task, wearing VR head-mounted display while either standing upright or lying supine on a bed. **c,** During the task, participants were navigating in a circular virtual arena, performing a spatial memory retrieval. At each trial: 1) the image of the target object was shown, 2) A participant navigated to the retrieved location, 3) As feedback, the target object appeared at its correct location to be collected.

## Results

### Participant’s physical BO significantly affects the experienced BO and navigation behavior

In order to assess the impact of physical BO on the experienced BO and navigational behavior in VR, we compared virtual navigation when our participants were supine vs. when they were standing without any avatar shown (Nobody conditions, Figure 1a blue & green). As predicted, we found a significant influence of physical BO on the experienced BO during the virtual navigation task in VR, as assessed through subjective questionnaire ratings and implicit behavioral measures. We found a significant effect of physical BO on questionnaire ratings pertaining to the experienced BO in VR (Figure 2a). Our participants reported significantly higher Q_Supine ratings (experience of being supine in VR; r = 0.783, p < 0.001) and lower Q_Standing ratings (experience of standing upright in VR; r = 0.593, p = 3.01e-03) when their physical BO was supine (i.e., Supine-Nobody condition) than standing (i.e., Standing-Nobody condition). These results indicate that an experienced BO in VR is influenced by a BO of the physical body in a direction that becomes congruent. Therefore, when participants’ physical BO was standing upright (i.e., Standing-Nobody), they felt as if they were standing in VR, rather than being supine (Q_Standing > Q_Supine; r = 0.84, p < 0.001). However, in the Supine-Nobody condition, Q_Supine and Q_Standing had equal ratings (i.e., did not differ; r = 0.093, p = 0.64), revealing an ambiguity in our participants’ experienced BO when their physical BO was supine and when they did not receive additional visual cues regarding BO in the virtual environment (i.e., Nobody condition). In addition, we found overall higher Q_Standing ratings compared to the Q_Supine (r = 0.566, p < 0.001), suggesting that our participants preferably experienced a standing BO in VR. The conflict score (see Methods) confirmed these findings, revealing higher conflict scores when participants were physically supine versus upright (r = 0.762, p < 0.001), possibly reflecting an incongruence between the participants’ physical BO (supine) and their preferably experienced BO in VR (upright).

**Figure 2.**
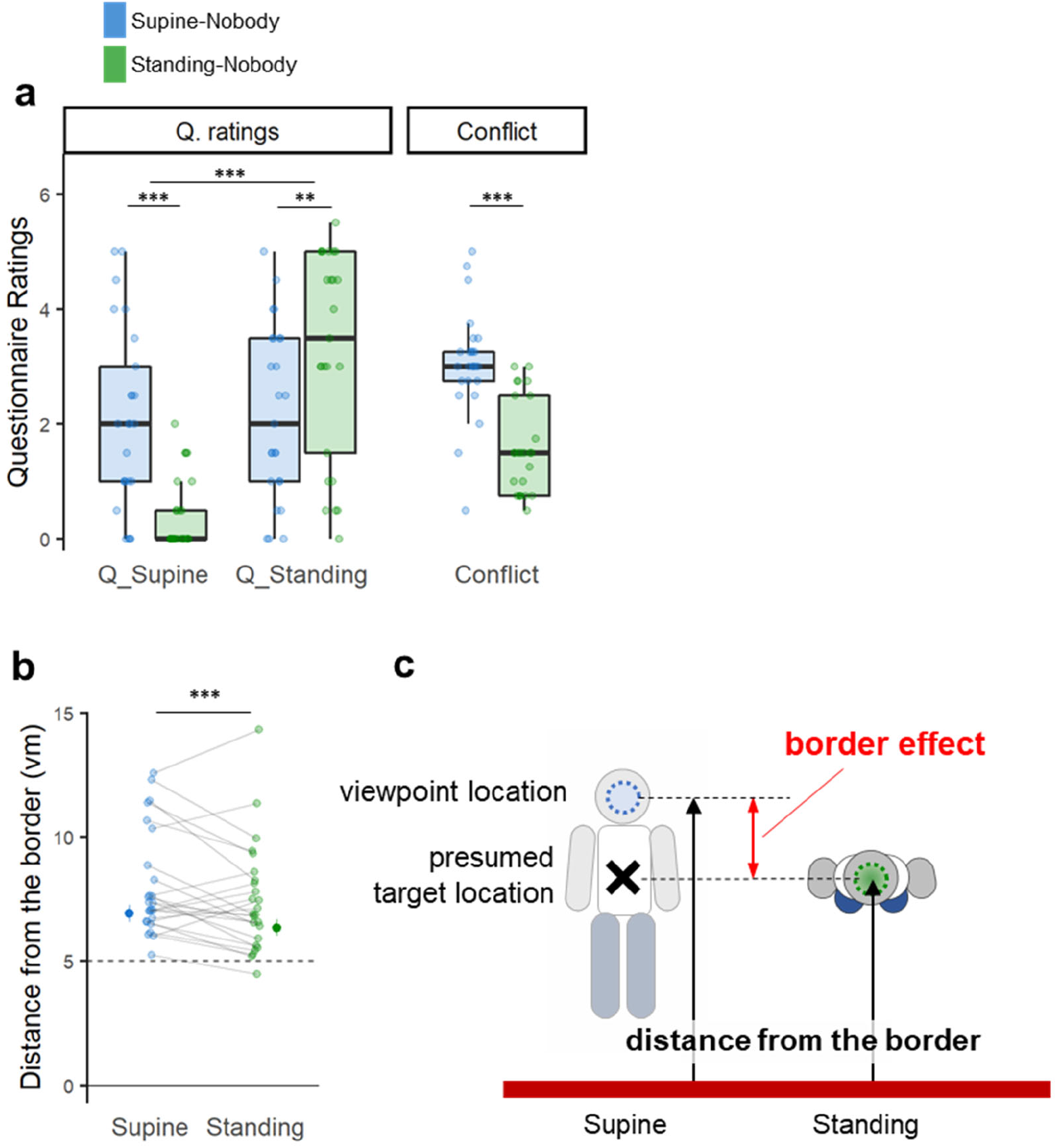
Effect of physical BO on experienced BO and on navigational behavior in VR. **a,** When participants were physically supine (blue), they had a stronger experience of being supine in VR while they felt less as if they were standing, compared to when they were actually standing upright. The conflict score indicates how much their experienced BO in VR conflicted with their physical BO. The results showed that the conflicts were significantly larger in the physically supine BO condition than in the standing BO condition. Also, overall, when compared regardless of their physical BO, participants reported a significantly stronger experience of being standing in VR, suggesting the standing BO as a presumed BO in VR. **b,** Participants’ reached location was farther away from the border when they were physically supine compared to when they were standing upright. **c,** A top-view schematic figure depicting the border effect. The changes in the distance from the border possibly reflected changes in self-location (or body boundaries) that are related to the change in the experienced BO in VR. Consequently, the distance from the border (calculated with respect to the viewpoint location) was greater when they were physically supine compared to when they were standing upright. **: 0.001 < = p < 0.01, ***: p < 0.001.

The significant influence of the physical BO on the experienced BO was further validated by implicit navigation behaviors during the task: the distance from the border when participants stopped at the end of each retrieval trial. Our participants stopped at a significantly larger distance from the border in Supine-Nobody condition compared to Standing-Nobody condition (mixed-effect regression; df = 1, F = 11.08, p < 0.001, n = 25; Figure 2b), as if they wanted to avoid that their legs would hit the border of the arena. We note that every task object was placed at a position equal distance (5 vm) away from the border and that participants were approaching the targets facing the border (majority of trials: 98.1%). Accordingly, we argue that the border effect (i.e., the larger/smaller distance from the border) reflects an implicit incorporation of the participant’s physical BO into navigation behavior (see (Alsmith and Longo, 2014; Moon et al., 2022)). Thus, even when not seeing a virtual agent during virtual navigation (as in Moon et al., 2022), supine participants stopped farther from the border, behaviorally corroborating the subjectively experienced supine BO (Figure 2c). Further analysis corroborated this association by showing that the distance from the border is significantly correlated with the strength of the participant’s subjective experience of being supine (Q_Supine) (df = 1, F = 19.08, p < 0.001, n = 25; Supplementary Fig. 5).

In addition, we also investigated the impact of the physical BO on other navigation performance measures (i.e., distance error, navigated distance, trial time). The impact of BO on these was assessed together with the effects with body view (also to check their possible interactions) and will be reported in a separate section (see the last section of the results).

### Effect of the virtual body on experienced BO and conflicts

We next analyzed the impact of the view of the BO-congruent avatar on experienced BO in VR. For this, we analyzed the two experimental conditions in which our participants were physically supine, comparing the condition showing the virtual scene without any avatar (i.e., Supine-Nobody condition) with the condition presenting the virtual scene and the supine virtual avatar (i.e., Supine-Body condition). In the latter condition, the avatar’s posture was congruent with the participants’ physical posture (supine). Assessing whether an avatar with a congruent BO (with respect to the participant’s physical BO) affects the experienced BO in VR, we show that when participants were physically supine and presented with an avatar (i.e., Supine-Body condition), they have an enhanced experience of being supine (Figure 3a; Q_Supine; r = 0.720, p < 0.001, n = 25) and a reduced experience of standing upright (Q_Standing; r = 0.595, p = 2.95e-03), compared to the condition without the avatar (i.e. Supine-Nobody condition). Furthermore, in Supine-Body condition, the ratings for Q_Supine were significantly higher than those for Q_Standing (Figure 3a; r = 0.759, p < 0.001), which was not the case in Supine-Nobody condition (r = 0.093, p = 0.64). The data, hence, demonstrate reduced ambiguity in experienced BO when the avatar was present. This is further confirmed by a lower conflict score in the condition with a body-congruent avatar compared to the condition with no avatar (r = 0.800, p < 0.001).

**Figure 3.**
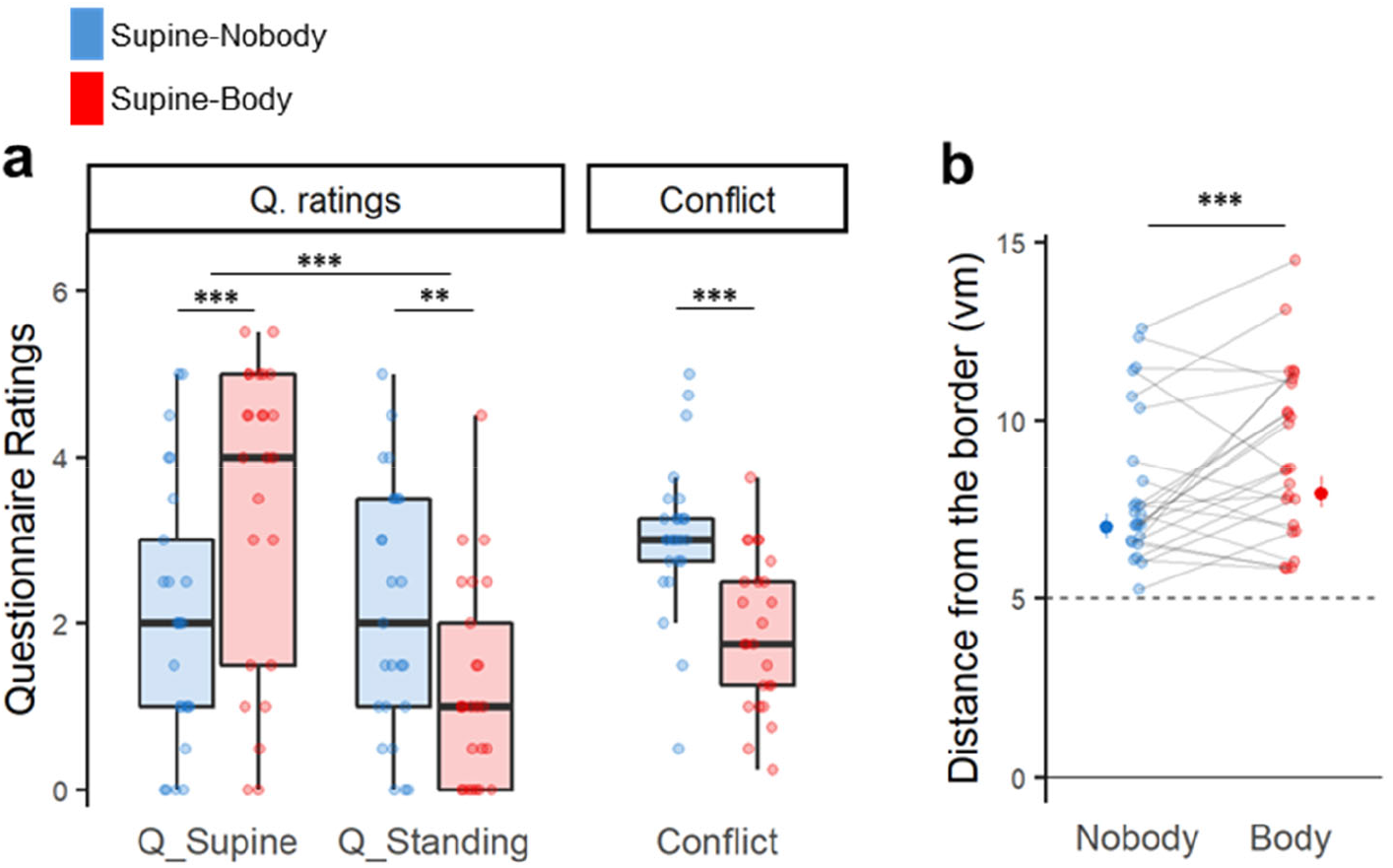
Experienced BO affected by the BO-congruent avatar. **a**, When the participants were physically supine, presenting an avatar with supine posture (i.e., Body condition) strengthened the subjective experience of being supine and reduced the experience of standing upright compared to the Nobody condition, where no avatar was shown. The conflict score confirms that showing the BO-congruent avatar significantly reduced the conflict of the BO. When compared regardless of the presence of the avatar, they felt more as if they were supine than standing, again highlighting the impact of the physical BO on the experienced BO in VR (Q_supine > Q_standing, overall). **b**, We also reproduced previous findings of a significant shift in the distance from the border between body and nobody conditions, suggesting that such drifts in self-location are associated with changes in the experienced BO in VR. **: 0.001 < = p < 0.01, ***: p < 0.001.

In addition, congruently with the questionnaire results, we found an even larger border effect (drift in self-location), with participants keeping larger distances from the border in Supine-Body condition compared to Supine-Nobody condition (mixed-effect regression; df = 1, F = 16.56, p < 0.001, n = 25; Fig 3b), confirming previous data by Moon et al. (2022). The association between the border effect and the experienced BO was confirmed by the significant correlation between the border distance and Q_Supine rating (df = 1, F = 19.08, p < 0.001, n = 25; Supplementary Fig. 5).

### Effect of physical BO and 1PP avatar on the navigation performances

Based on the data from four of our experimental conditions, we assessed the impact of both physical BO and presenting a supine virtual avatar on navigation performance. Either of those effects and their possible interaction were simultaneously taken into account through a dedicated mixed-effect model for each behavioral parameter. Through these analyses, we found a significant impact of physical BO on trial time: the time participants spent per retrieval trial was significantly shorter when they were physically standing upright versus supine (mixed-effect regression; df = 1, F = 32. 90, p < 0.001, n = 25; Supplementary Figure 4c). However, we did not observe any significant influence of physical BO on spatial memory accuracy (i.e., distance error) and navigated distance per trial. On the other hand, seeing a virtual agent while navigating (i.e., Body condition) significantly reduced navigated distance compared to the conditions without an avatar (mixed-effect regression; df = 1, F = 25. 95, p < 0.001; Supplementary Figure 4d), while it did not affect the other navigation performance measures (i.e., distance error, trial time). This suggests more efficient navigation (i.e., less distance traveled to reach the same location) in the avatar condition. Notably, we did not find significant interactions between physical BO and avatar conditions on any of the navigation performance measures. Overall, these data suggest that the supine physical BO (i.e., BO in MRI) had significant negative impacts on virtual navigation performance (i.e., trial time) and that seeing a virtual agent can improve spatial navigation performance in VR (i.e., navigated distance per trial).

## Discussion

In this study, we demonstrate that physical BO (standing or supine participants) modulated experienced BO and navigation behavior in VR. Our data also show that a standing BO is preferred during spatial navigation, because participants were more likely to experience a standing BO than a supine BO and because their navigation performance was better when they were standing upright (i.e., shorter navigation time). We also corroborate previous reports by demonstrating that showing a virtual avatar in a supine position reduces the conflict between supine physical BO and preferred BO (i.e., standing) and improves spatial navigation (i.e., shorter navigation path). Because many navigation-related neuroimaging studies are done in the MRI scanner(Doeller et al., 2010; Kunz et al., 2015; Horner et al., 2016; Stangl et al., 2018; Bierbrauer et al., 2020; Moon et al., 2022), where physical BO is constrained to the supine BO(Taube et al., 2013; Steel et al., 2021), our study provides important information that should be considered when spatial navigation studies in the scanner are designed or interpreted.

First, we demonstrated that participants’ physical BO modulated their experienced BO in VR as measured by subjective questionnaire ratings and by the border effect. Concerning BO ratings, we found that when participants navigated the virtual environment without seeing an avatar, their experienced BO was congruent with their physical BO: Standing BO ratings increased and supine BO ratings decreased when they were physically standing upright, and vice versa. These subjective data were corroborated by the border effect in navigation behavior (i.e., the distance from the navigation arena’s border during retrieval trials). When navigating to the target location, our participants kept a larger distance from the border while their physical body was in supine versus standing BO (Figure2-b,c). We note that both conditions were visually identical (no avatar in either condition) and differed only in physical BO. Thus, the border effect cannot result from visual differences between conditions (i.e., with or without an avatar), but rather by the different physical BO, associated with differences in experienced BO: the space occupied by the body expands forward in the supine BO (also see (Alsmith and Longo, 2014; Moon et al., 2022)). The finding of a positive correlation between the distance from the border with the strength of the participant’s subjective experience of being supine confirms this interpretation (Supplementary Figure 5). Accordingly, we argue that the border effect is an implicit change in navigational behavior when participants are in supine physical BO vs. standing. Importantly, this navigational parameter provides an objective and repeated proxy of one’s experienced BO, eliminating potential biases that may arise from explicit questionnaires. This effect is of direct relevance for spatial navigation studies performed during fMRI (i.e.,(Doeller et al., 2010; Kunz et al., 2015; Stangl et al., 2018; Bierbrauer et al., 2020)) where supine BO is unavoidable. Apart from altering navigation behavior, the supine BO may also affect place/grid-cells-related brain activity (i.e., GCLR signals in entorhinal cortex; (Doeller et al., 2010; Jacobs et al., 2013; Nadasdy et al., 2017; Maidenbaum et al., 2018; Moon et al., 2022)) or the activation of other regions involved in spatial navigation such as retrosplenial cortex(Vann et al., 2009; Mitchell et al., 2018; Bierbrauer et al., 2020; Alexander et al., 2023). The influence of physical BO on the experienced BO suggest that a virtual agent (even when it is invisible in Nobody condition) and the participant’s body are functionally linked to each other during virtual navigation. The association could be mediated by the intrinsic signals from the physical body (e.g., vestibular and proprioceptive signals)(Pfeiffer et al., 2014; Lenggenhager and Lopez, 2015; Park and Blanke, 2019). Thus, direct changes or disruptions to the bodily inputs could influence both the navigation experience (BO) and related navigation behaviors (i.e., border effect). This hypothesis needs further investigation in future studies using experimental manipulation of those bodily signals, such as electrical vestibular stimulation (Sluydts et al., 2020).

Secondly, our data show that standing is the preferred experienced BO during virtual navigation. We found that a conflict between the physical BO and the experienced BO in VR (as measured by the conflict score) was larger when our participants were physically supine versus standing upright. Thus, when they were physically standing upright, they felt as if they also were standing in the virtual arena during navigation (supported by bodily signals). In contrast, when they were in supine BO in the real world, their reported experience of BO in the virtual arena was more ambiguous (between supine and standing). Considering the influence of the supine physical BO we discussed above, the observed ambiguity could be the result of incongruence between the preferred BO in VR (i.e., standing upright) and the supine physical BO (arguably, mediated by vestibular and proprioceptive cues from the supine body). We argue that the preferred standing BO during virtual navigation most likely originates from our daily experience of upright position during physical navigation. Human brain mechanisms of spatial navigation could have adapted to the evolutionary change in BO and been optimized for navigation in standing upright BO (also based on body structure, sensory system, and lifestyle distinguished from other animals (Ekstrom, 2015)). This was also indirectly supported by influences of the physical BO and experienced BO on spatial navigation performance in VR. We observed a decrease in time to retrieve and navigate to the target when participants’ physical BO was standing (Supplementary Figure 4c), suggesting that standing BO (i.e., preferred BO during navigation as suggested above) facilitates some spatial navigation processes. Alternatively, this effect could also be linked to changes in space perception (e.g., visual vertical judgment; which is possibly related to the prediction of the destination ahead of straight navigation) that have been reported to be better in the upright position (compared to the supine position), by the contribution of vestibular gravitational signals (Lopez et al., 2008; Lopez and Blanke, 2010). Size or distance perceptions have been reported to be better while upright than supine: supine BO makes the size or the distance more underestimated(Kim et al., 2022). Alternatively, although the VR scene presented in the HMD was the same, our participants might have experienced it from an elevated perspective (i.e., as if they were looking down; which is a more likely situation while standing than supine) when they were physically upright but not when they were supine. An elevated perspective has been associated with faster response times in visuospatial tasks (vs. eye-level or lowered perspective) (Schwabe et al., 2009). Indeed, a previous study by Ionta and colleagues (Ionta et al., 2011) reported that BO in a virtual space experienced by participants in the MRI scanner (i.e., looking-up vs. looking-down) could be altered by multisensory bodily signals while keeping the visual input constant. Altogether, the present findings suggest that the human navigation system “has a preference” for navigation in the upright BO, and that - although virtual navigation can simulate natural navigation (as if upright) with some extent of ambiguity - it may compromise the navigation performance in supine BO (as in MRI).

Finally, we show that also the presentation of a supine avatar during virtual navigation influences both experienced BO and navigation behavior in VR. Concerning experienced BO in VR, we report that the avatar in the supine position was associated with a stronger sensation of being in supine BO in VR. This finding is compatible with an important role of bodily multisensory cues in virtual navigation experience. It should be stressed that this rather simple experimental manipulation (i.e., showing a supine avatar during virtual navigation) significantly reduced the conflict between the experienced BO and the physical BO when participants were lying supine (which is the adopted posture in MRI acquisitions), as highlighted by the lower conflict score in Supine-Body condition compared to Supine-Nobody condition. As discussed above, participants’ experienced BO in the Supine-Nobody condition (i.e., the typical condition in MRI) is experienced as ambiguous, arguably due to the incongruency between preferred BO and the influence of supine physical BO. However, our results suggest that the addition of a seen avatar reduces this ambiguity and the related sensory-experiential conflict (i.e., conflict score). Importantly, we show that these subjective BO changes were reflected in the border effect, reproducing previous results in the MRI scanner (Moon et al., 2022). Of note, this border effect, as induced by a supine avatar (Supine-Body condition), further increased the distance from the border compared to Supine-Nobody condition, where we observed the border effect induced by supine physical BO when compared to Standing-Nobody. Thus, the border effect further increased when the subjective experience of being supine in VR was enhanced by (1) supine physical BO and again by (2) the presence of a supine avatar (Standing-Nobody < Supine-Nobody < Supine-Body). This finding was further supported by the significant correlation between the border effects and Q_supine ratings, and vice versa (Supplementary Fig. 5b). Moreover, seeing a supine avatar also reduced the navigated distance per trial without increasing retrieval errors, suggesting improved spatial navigation in the conditions with an avatar (Supplementary Figure 4d) as was in the similar study in MRI (Moon et al., 2022). This is again of relevance to human navigation studies using virtual navigation paradigms, and in particular to those paradigms employing MRI (Taube et al., 2013; Steel et al., 2021). Overall, our data suggest that body-related cues (e.g., view of a body-posture-congruent avatar) as well as signals from the body systematically evoke conflicts to different degrees depending on the actual conditions; this knowledge should be used to improve and better understand experienced BO and performance in virtual navigation, especially when a supine BO is necessary as in the MRI studies (Doeller et al., 2008; Doeller et al., 2010; Taube et al., 2013; Stangl et al., 2018; Bierbrauer et al., 2020; Moon et al., 2022).

Collectively, our results highlight the importance of the physical BO and the relevant visual cue (i.e., BO-indicating body view) on virtual navigation by showing their influence on both subjectively experienced BO and the navigation behaviors in VR. Through this study, we confirm that standing BO is preferred during navigation with solid evidence, which has been considered so without much evidence. By comparing the conventional condition in MRI with others (including the preferred condition) across a range of aspects, our data provide a more fine-grained understanding of the potential, but often overlooked, effects that could be caused by the constraint in MRI. Importantly, the present data show that the addition of seen avatar can be a simple but powerful method to overcome the limitation of navigation in MRI, resolving the ambiguity of the experience of BO and improving navigation performance (i.e., Moon et al., 2022). Our results strongly suggest that multisensory body-related aspects should be cautiously considered in the experimental design of any virtual navigation study. Finally, we propose the border effect as a robust measure that allows assessing the potential biases and confounds in experienced BO during virtual navigation, and thereby helps control them.

## Materials and Methods

### Participants

Twenty-five healthy participants (11 males and 14 females; mean age 25.7±1.91) participated in the study. They gave informed consent following the institutional guidelines (IRB #: GE 15-273) and the Declaration of Helsinki (2013). They were neither aware of the purpose of the study nor had a history of psychological disorders. They were right-handed with normal or corrected-to-normal vision. They were recruited from the general population through the online recruitment system and received monetary compensation according to the contributed time (CHF20/hour). The number of participants, 25, was chosen to match the sample size of the previous study using a similar spatial navigation paradigm (Moon et al., 2022). Participants who quit the experiment (four; three females) due to severe motion sickness were already excluded from the dataset and are not counted in the sample size.

### Virtual Reality (VR) spatial navigation task with Head-mounted display (HMD)

The spatial navigation task used in the study was implemented with Unity Engine (Unity Technologies) by adapting the paradigm from the previous study (Moon et al., 2022). The participants wore the head-mounted display (HMD; Oculus Rift S, Oculus) and used a gaming joystick (Extreme 3D Pro, Logitech) to perform the task in the circular virtual arena. Distal landmarks were placed outside of the task arena, providing orientation cues to the participants. Each session of the task began with an encoding phase, during which participants navigated the arena and encoded the positions of the three task objects placed within it. Next, they performed the recall task composed of 14 trials. In each recall trial, the target object was shown for two seconds (i.e., Cue phase) and participants had to recall its original location and navigate to there (i.e., Retrieval phase). Finally, the object reappeared in its correct position providing feedback to the participants, and had to be collected again in order to trigger the start of the next trial. In the last trial, a threatening scenario (i.e., a virtual knife approaching from the sky toward either the avatar or the space where the avatar should have been placed in Nobody condition) was presented to the participants. This was to provide an additional measure (i.e., response to the threat) of self-identification and self-projection of participants, the association between the virtual agent in VR and the sense of self.

The navigation task was performed in four different conditions obtained from the combination of the avatar’s presence/absence (i.e., Body vs. Nobody) and the physical BO during the task (i.e., Supine vs. Standing BO) (Figure 1). Of note, while the two manipulations generate four conditions, our design is not a two-by-two design. In fact, the avatar was only ever presented in a supine position (congruent to Supine-Body condition), while no standing avatar was shown in the Nobody conditions. Therefore, we had no congruent avatar condition for the standing BO. Supine-Body condition was adopted to assess the effect of a body-congruent avatar as was Moon et al. (2022), while Standing-Body condition was adopted to assess how incongruency between the participant’s physical BO and the avatar’s BO modulates subjectively experienced BO and spatial navigation performance.

The whole experiment was separated into two blocks. In each block, participants went through all four conditions presented in different orders. To avoid fatigue due to prolonged standing, physical BOs (i.e., standing and supine) were interleaved with each other within a block. The order of the conditions was pseudo-randomized and counterbalanced between participants. In total, each participant performed eight sessions of the navigation task (twice per condition) and answered the questionnaires at the end of each session (see Questionnaire section for the details).

### Physical BO during the task

To assess the impact of the physical BO on the experienced BO during the virtual navigation, we asked participants to perform the experiment in two different physical BO: in the supine BO condition, participants lay down on the bed with the joystick positioned on their right-hand side (Figure 1b-left); in the standing BO condition (Figure 1b-right) they stood upright and the joystick was positioned on a height-adjustable table on their right side. The height of the bed during the supine condition was set to approximately match the height of the participant’s upper trunk when they stood up.

### Virtual avatar during the task

A virtual avatar in a supine position was presented in the Body condition of the task (note that the avatar posture was always supine regardless of the BO condition). The movements of the avatar’s right hand were programmed to match the participants’ hand movements while controlling the joystick, providing a visuo-motor congruency. In the supine BO condition with the avatar (i.e., Supine-Body condition), such visuo-motor congruency, together with the visuo-proprioceptive congruency of the physical BO and the virtual avatar’s BO, was expected to induce a higher illusory self-identification with the avatar during navigation. By contrast, in the standing BO condition with the avatar (i.e., Standing-Body condition) this visuo-proprioceptive congruency was not met as the physical BO was incongruent with the avatar’s BO. Therefore, we expected lower self-identification with the avatar under this condition (Sanchez-Vives et al., 2010; Slater et al., 2010; Walsh et al., 2011; Kokkinara and Slater, 2014; Blanke et al., 2015).

### Questionnaire

At the end of each session, participants were asked to rate their agreement with seven statements using a Likert scale ranging from 0 (strongly disagree) to 6 (strongly agree). All the items are listed in Supplementary Table 1. The order of the statements was shuffled at every session and participants rated them autonomously with the joystick. Q1, Q2, and Q3 were aimed at assessing the bodily self-consciousness of the participants. Q6 and Q7 were designed to probe their experienced BO during the virtual navigation. The conflict between the physical BO and experienced BO was quantified as the conflict score, which was calculated with the participants’ physical BO and the ratings of Q6 and Q7 by the following formula:

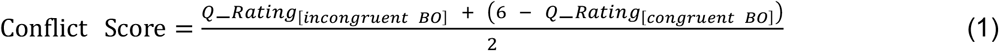

For instance, when a participant was physically supine, the conflict score was calculated as (Q_Standing + (6 - Q_Supine))/2: the higher when their experienced BO was opposed to the physical BO. Q4 was aimed to capture the cyber-sickness during the task, while Q5 served as a general control question. We shortly debriefed the participant once the entire experiment was completed, to ensure the integrity of their autonomous responses and also to record their spontaneous subjective reports.

### Statistical analysis

All the behavioral and questionnaire data were analyzed using R (v4.1.2 for Windows, https://www.r-project.org/) and RStudio (v2021.09.01, http://www.rstudio.com). The differences in the questionnaire ratings and conflict scores between conditions were assessed with a paired Wilcoxon signed-rank test. For the behavioral parameters recorded at each trial (i.e., distance errors, navigation trace length and time, distance from the border), we performed mixed-effects regressions (lme4, v1.1-18-1) with a fixed effect of condition and random intercepts for individual participants to assess statistical significance. Random slopes were assumed as far as the model did not fail to converge. We also examined the correlations between parameters using mixed-effect regression models. The distribution of each dependent variable was considered in the mixed-effect modeling of the variable, following the previous study using a similar task (Moon et al., 2022).

## Acknowledgments

This work was supported by the Korea Institute of Science and Technology (KIST) Institutional Program (2E32341), and the Bertarelli Foundation to H.-J.M. O.B. is supported by the Swiss National Science Foundation (No. 320030_188798) and by the Bertarelli Foundation. Additional support was provided by the Fondation Campus Biotech Geneva (FCBG)—a foundation of the Swiss Federal Institute of Technology Lausanne (EPFL), the University of Geneva (UniGe), and the Hôpitaux Universitaires de Genève (HUG), the Institute of Translational Molecular Imaging (ITMI).

## Author Contributions

Conceptualization, H.-J.M., H.-P.W., E.D.F., and O.B.; Methodology, H.-J.M. and H.-P. *W*.; Software, H.-J.M.; Formal Analysis, H.-J.M.; Investigation, H.-J.M., H.-P. *W*., and E.D.F.; Writing – Original Draft, H.-J.M.; Writing – Review & Editing, E.D.F., H.-P. *W*., and O.B.; Resources, H.-J.M.; Funding Acquisition, O.B.; Supervision, O.B.

## Competing Interests

The authors declare no competing interests.

## Data and code availability

The data that support the findings of this study and the analysis code are available in ‘github’ through the following link [https://github.com/JuneHMoon/Moon_et_al_BO_during_VR.git]

## Notes

### Competing Interest Statement

The authors have declared no competing interest.

