## Supplementary Information for "Real-world body orientation impacts virtual navigation experience and performance"

### Supplementary Text

##### Standing-Body condition: The combined effect of standing physical BO and the supine avatar on the experienced BO

In a fourth condition, the Standing-Body condition (Figure 1a-yellow), we combined the standing physical BO with the visualization of a supine and motion-mimicking avatar in first-person perspective (1PP). We compared this condition with Supine-Nobody condition (Figure 1a-blue), corresponding to the classical condition used in most MRI studies on navigation, to assess the combined effects of standing BO and presenting a supine avatar, and their possible interaction (due to any specific effect of BO-congruency between the physical body and the avatar). An effect of the BO-congruent avatar on self-identification was confirmed by the significantly higher ratings in the questionnaires in Supine-Body condition versus Supine-Nobody condition (Q_Self-id: $r = 0.653$, $p = 1.10e-03$, $n = 25$; Q_Threat: $r = 0.538$, $p = 7.11e-03$; Supplementary Fig. 2). In contrast, when one's physical BO was incongruent with the avatar's BO (i.e., Standing-Body condition), neither Q_Self-id or Q_Threat was significantly different from Supine-Nobody condition (Supplementary Fig. 3).

By design, the effect of standing physical BO, enhancing the feeling of standing upright in VR, and the effect of a supine 1PP avatar, strengthening the feeling of being supine, were expected to go in opposite directions. We found that, despite seeing a supine avatar from 1PP, participants reported higher ratings for Q_Standing than Q_Supine in Standing-Body condition ($r = 0.45$, $p = 0.015$; Supplementary Figure 3b). In addition, participants reported significantly weaker feelings of being supine in Standing-Body condition compared to Supine-Nobody condition ($r = 0.44$, $p = 0.028$, $n = 25$). These results again suggest a strong influence of physical BO that could override the avatar-induced effect on the subjective experience of BO in VR.

### Supplementary Table

#### Supplementary Table 1

**Supplementary Table 1. Questionnaire**

| Questionnaire | | |
|---|---|---|
| Q1 | Self-identification | I felt as if what I saw in the middle of the scene was my body. |
| Q2 | Threat | I felt as if the threat(knife) was toward me. |
| Q3 | Presence | I felt as if I was located in the virtual environment. |
| Q4 | Cyber-Sickness | I felt dizzy. |
| Q5 | Control | I felt as if I had 3 bodies. |
| Q6 | Supine | I felt as if I was supine in the virtual environment. |
| Q7 | Standing | I felt as if I was standing in the virtual environment. |

### Supplementary FIGURES

#### Supplementary Figure 1

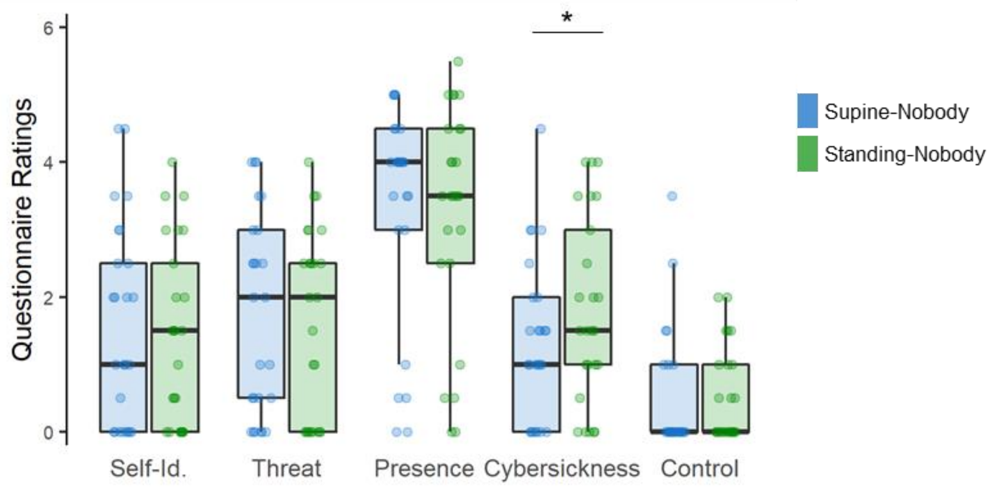


**Fig. S1, related to Fig. 2, Questionnaire ratings of the Supine-Nobody and Standing-Nobody conditions.** Participants' ratings regarding self-identification (Self-Id), response to threat (Threat), presence, and control did not differ significantly whether they were physically supine or standing. They reported higher ratings for cybersickness in the Standing-Nobody condition than in the Supine-Nobody condition. *: $p < 0.05$

#### Supplementary Figure 2

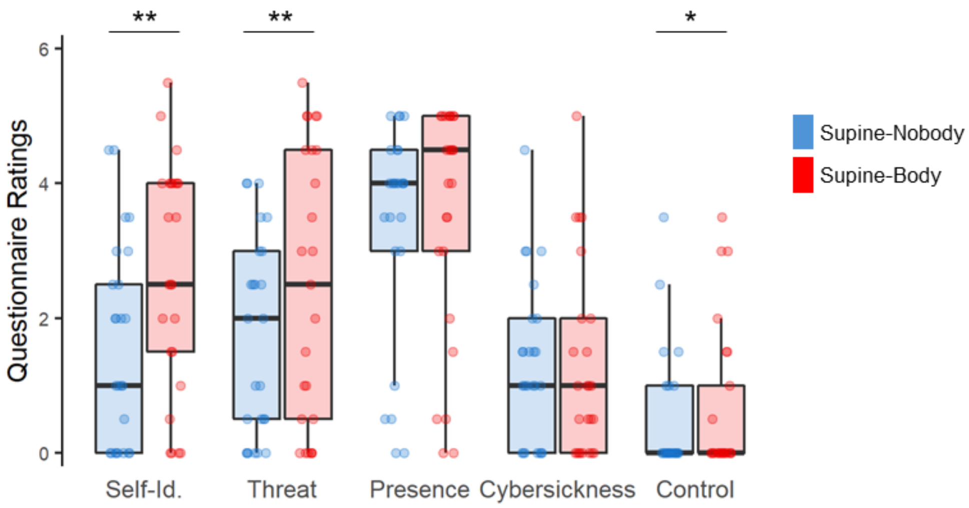


**Fig. S2, related to Fig. 3, Questionnaire ratings of the Supine-Nobody and Supine-Body conditions.** Participants' ratings regarding self-identification (Self-Id) and response to threat (Threat) were significantly higher in the condition with posture- and motion-congruent avatar (i.e., Supine-Body condition) than the condition without an avatar (i.e., Supine-Nobody). While ratings for presence and cybersickness did not differ significantly between the two conditions, we found a difference in the control questionnaire ratings: higher in the Supine-Body than the Supine-Nobody. *: $p < 0.05$, ** : $0.001 <= p < 0.01$

#### Supplementary Figure 3

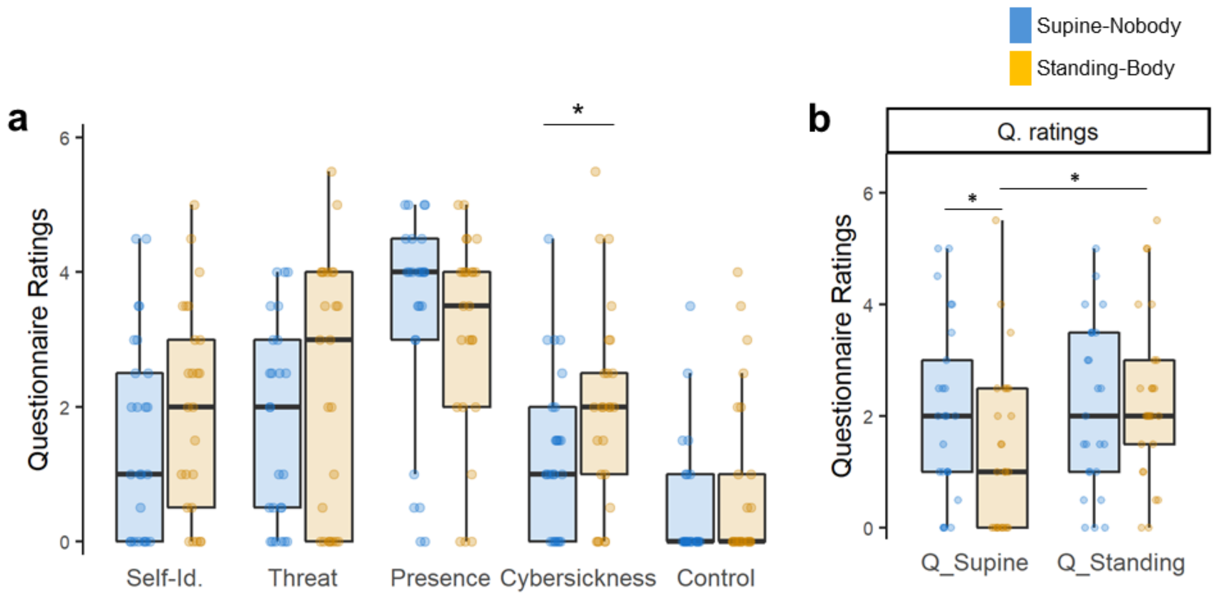


**Fig. S3, Questionnaire ratings of the Supine-Nobody and Standing-Body conditions. a,** Participants' ratings regarding self-identification (Self-Id) and response to threat (Threat) were not significantly different between the Supine-Nobody and Standing-Body conditions. Although the standing-Body condition also showed a supine motion-mimicking avatar, as was in the Supine-Body condition, the avatar-induced bodily self-consciousness changes were not significant when the posture of the avatar was incongruent with the physical body posture. We found higher ratings for cybersickness in the Standing-Body condition than in the Supine-Nobody condition. **b,** The Standing-Body condition led to significantly higher ratings for feeling standing compared to feeling supine (the two yellow boxes), in contrast, the two ratings did not significantly differ from each other in the Supine-Nobody condition (blue boxes). Also, participants rated significantly lower for feeling supine (Q_Supine) in the Standing-Body condition compared to the Supine-Nobody condition. These results suggest that participants' experienced BO was more affected by the physical standing BO than by the presence of the supine avatar. *: $p < 0.05$

#### Supplementary Figure 4

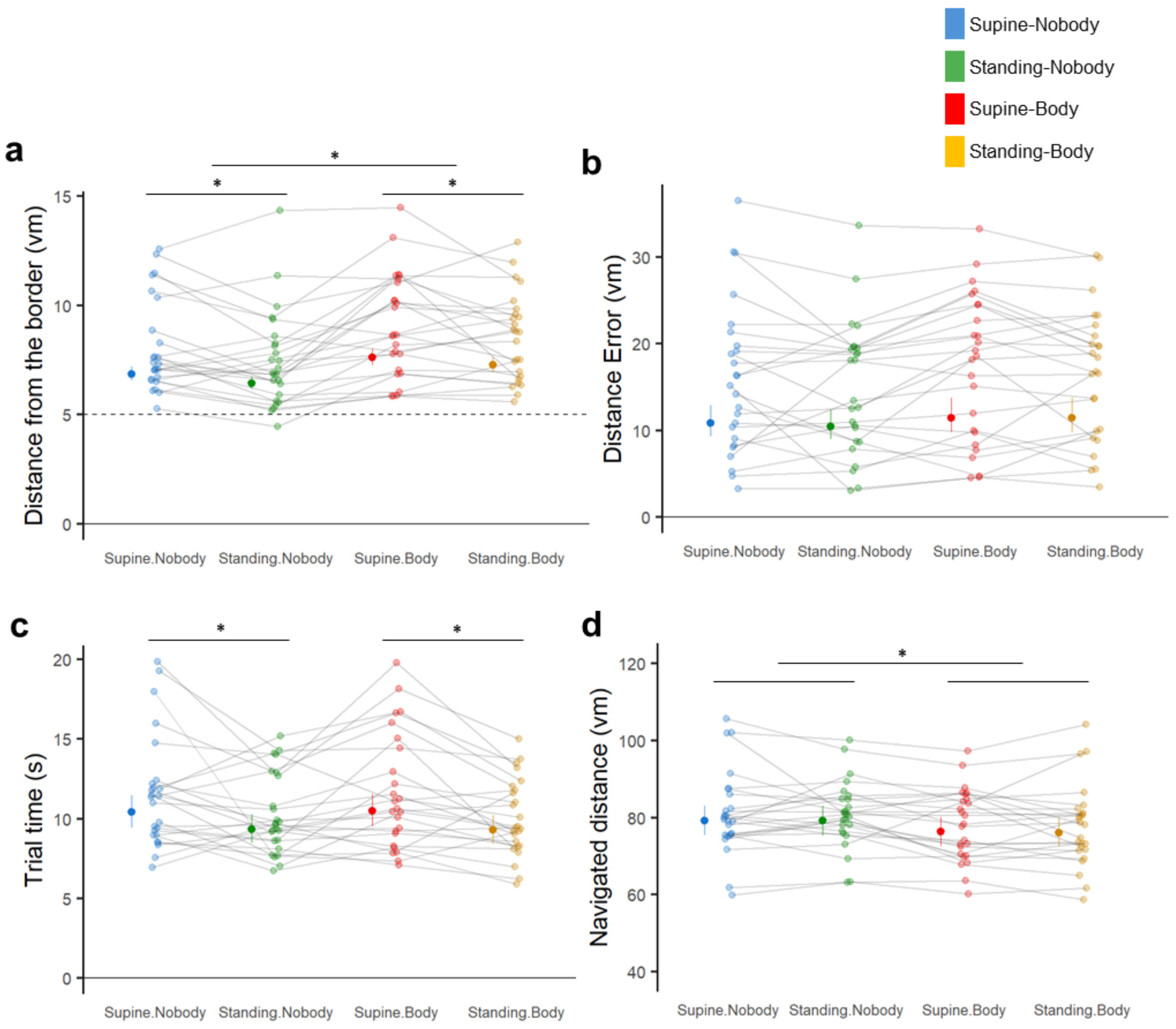


**Fig. S4, Navigation parameters of four experimental conditions.**
**a,** Distance from the border, which arguably reflects experienced BO in VR, was significantly larger when participants were physically supine than standing, and also when a supine avatar was presented in a first-person viewpoint position than no avatar was shown. **b,** Distance Error, indexing spatial navigation precision, did not significantly differ among the four conditions. **c,** Participants spent significantly less time in the retrieval phase when they were physically standing compared to when they were physically supine, regardless of the presence of the avatar. **d,** The navigated distances were significantly reduced in the Body conditions (i.e., Supine-Body & Standing-Body) compared to the Nobody conditions (i.e., Supine-Nobody & Standing-Nobody). No significant interactions were found for all four parameters. A dedicated mixed-effect model was used for statistical assessments of each parameter. *: $p < 0.05$

#### Supplementary Figure 5

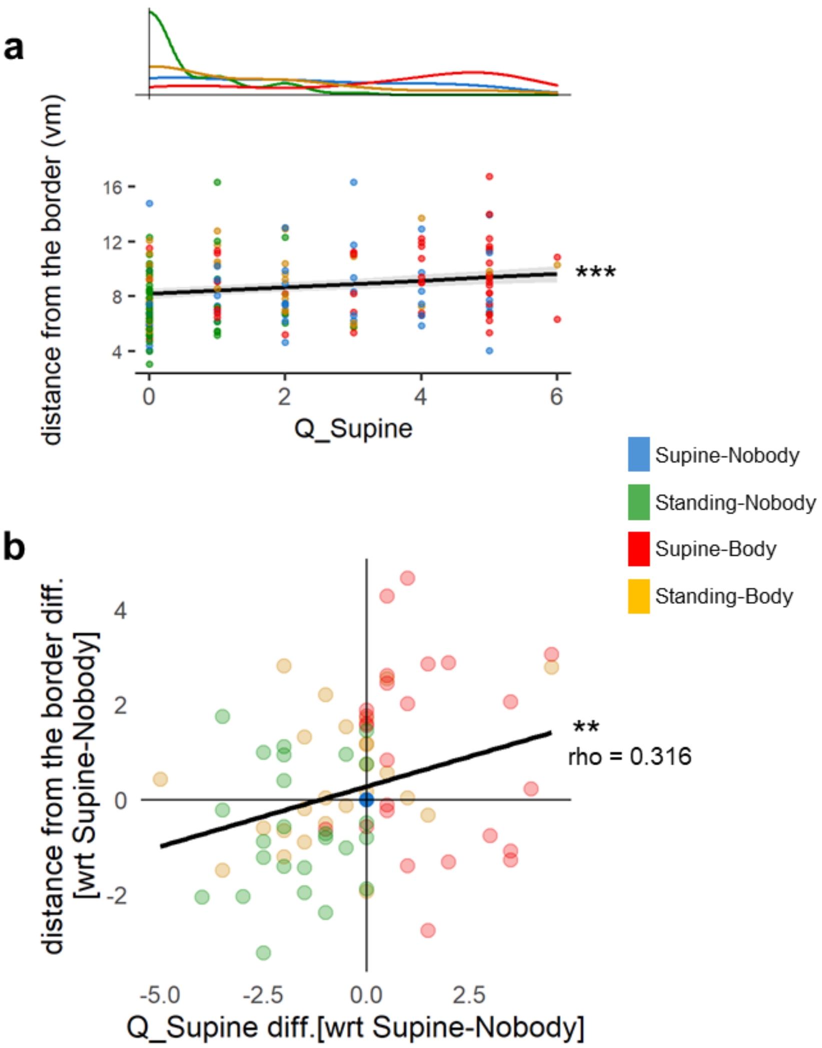


**Fig. S5, Distance from the border serves as an objective behavioral measure of experienced BO (i.e., supine position) in VR. a,** Distance from the border data were significantly correlated with ratings of Q_Supine (df = 1, F = 19.08, $p < 0.001$, n = 25). A mixed-effect model was used to assess their relationship. **b,** Their relationship at the within-subject level was further assessed through in-depth analysis. The distance from the border and Q_supine data were re-calculated and plotted with respect to the Supine-Nobody condition (i.e., scanner condition). We found that a change in the distance from the border of a subject in a condition was significantly associated with the change in the Q_supine rating of the subject in the condition. **: $0.001 <= p < 0.01$, ***: $p < 0.001$
